# Incomplete annotation of disease-associated genes is limiting our understanding of Mendelian and complex neurogenetic disorders

**DOI:** 10.1101/499103

**Authors:** David Zhang, Sebastian Guelfi, Sonia Garcia Ruiz, Beatrice Costa, Regina H. Reynolds, Karishma D’Sa, Wenfei Liu, Thomas Courtin, Amy Peterson, Andrew E. Jaffe, John Hardy, Juan Botia, Leonardo Collado-Torres, Mina Ryten

## Abstract

There is growing evidence to suggest that human gene annotation remains incomplete, with a disproportionate impact on the brain transcriptome. We used RNA-sequencing data from GTEx to detect novel transcription in an annotation-agnostic manner across 13 human brain regions and 28 human tissues. We found that genes highly expressed in brain are significantly more likely to be re-annotated, as are genes associated with Mendelian and complex neurodegenerative disorders. We improved the annotation of 63% of known OMIM-morbid genes and 65% of those with a neurological phenotype. We determined that novel transcribed regions, particularly those identified in brain, tend to be poorly conserved across mammals but are significantly depleted for genetic variation within humans. As exemplified by *SNCA*, we explored the implications of re-annotation for Mendelian and complex Parkinson’s disease. We validated in silico and experimentally a novel, brain-specific, potentially protein-coding exon of *SNCA*. We release our findings as tissue-specific transcriptomes in BED format and via vizER: http://rytenlab.com/browser/app/vizER. Together these resources will facilitate basic genomics research with the greatest impact on neurogenetics.

## Introduction

Genetic and transcriptomic studies are fundamentally reliant on accurate and complete human gene annotation. Gene definitions (namely genic coordinates and the isoforms/exons of which they are composed) are required for the quantification of expression or splicing from RNA-sequencing experiments, interpretation of significant genome-wide association studies (GWAS) signals and variant interpretation from genetic tests. As our understanding of transcriptomic complexity improves it is apparent that existing annotation remains incomplete even amongst known genes. Comparison of different gene annotation databases reveals that over 17,000 Ensembl genes fall into intronic or intergenic regions according to the AceView database and predictably, the choice of reference annotation greatly influences the output of variant interpretation software such as VEP and ANNOVAR^1,2^. Thus, incomplete annotation may cause pathogenic variants to be overlooked within exonic regions that are yet to be annotated and limit our understanding of risk loci.

Importantly, the impact of incomplete annotation of the transcriptome may not be evenly distributed across all types of tissues or cells and there is reason to believe that improvements to gene annotation may have a disproportionate impact on the understanding of Mendelian and complex neurological diseases. This view is supported by several analyses of bulk RNA-sequencing data derived from human brain tissues, which have discovered transcription originating from intronic or intergenic regions (henceforth termed novel)^3–5^. In particular, Jaffe and colleagues found that as much as 41% of transcription in the human frontal cortex was novel. Furthermore, it is becoming increasingly clear that RNA processing is highly complex in the human central nervous system due to the expression of long genes and the large diversity of cell types present^6,7^. In combination these factors could result in rare yet important transcripts being overlooked.

In this study, we address this issue by leveraging publicly available transcriptomic data available through the Genotype-Tissue Expression Consortium (GTEx) to improve the annotation of genes across the genome. We define transcription in an annotation-agnostic manner using RNA-sequencing data from 13 regions of the human central nervous system and compare this to definitions generated from a further 28 GTEx non-brain tissues. While we discover novel transcription to be widespread across all tissues, it is most prevalent in human brain. We provide evidence to suggest that the additional annotations we generate are likely to be functionally important on the basis of the tissue and cell-type specificity of novel expressed regions (ERs), the significant depletion of genetic variation amongst humans within ERs and their protein coding potential. Finally, by combining novel expressed regions (ERs) with split read data, defined as reads that have a gapped alignment to the genome, we link these regions to known genes associated with Mendelian and complex neurodegenerative and neurospsychiatric disorders. We release our findings as tissue-specific transcriptomes in a BED format and in an online platform vizER (http://www.rytenlab.com/browser/app/vizER), which allows individual genes to be queried and visualised. Overall, we improve the annotation of 1929 (63%) OMIM genes and a further 317 genes associated with complex neurodegenerative and neuropsychiatric disease. We anticipate that this will lead to improvements in diagnostic yield from whole genome sequencing (WGS) and the understanding of neurogenetic disorders.

## Results

### Optimising the annotation-agnostic detection of transcription using known exons

Pervasive transcription of the human genome, the presence of pre-mRNA even within polyA-selected RNA-sequencing libraries and variability in read depth complicates the identification of novel exons and transcripts using RNA-sequencing data^8,9^. With this in mind, we identified a set of exons with the most reliable and accurate boundaries (namely all exons from Ensembl v92 that did not overlap with any other exon^10^) and then used this exon set to calibrate the detection of transcription from 41 GTEx tissues^11^. We used the well-established tool, *derfinder*, to perform this analysis^12^. However, we noted that while *derfinder* enables the detection of continuous blocks of transcribed bases termed expressed regions (ERs) in an annotation-agnostic manner, the mean coverage cut-off (MCC) applied to determine transcribed bases is difficult to define and variability in read depth even across an individual exon can result in false segmentation of blocks of expressed sequence. Therefore, in order to improve our analysis and define ERs more accurately, we applied *derfinder*, but with the inclusion of an additional parameter we term the max region gap (MRG), which merges adjacent ERs (see detailed Methods). Next, we sought to identify the optimal values for MCC and MRG using our learning set of known, non-overlapping exons.

This process involved generating 506 transcriptome definitions for each tissue using unique pairs of MRCs and MRGs, resulting in a total of 20,746 transcriptome definitions across all 41 tissues. For each of the 20,746 transcriptome definitions, all ERs that intersected non-overlapping exons were extracted and the absolute difference between the ER definition and the corresponding exon boundaries, termed the exon delta, was calculated (**Figure 1a**). We summarised the exon delta for each transcriptome using two metrics, the median exon delta and the number of ERs with exon delta equal to 0. The median exon delta represents the overall accuracy of all ER definitions, whereas, the number of ERs with exon delta equal to 0 indicates the extent to which ER definitions precisely match overlapping exon boundaries. The MCC and MRG pair that generated the transcriptome with the lowest median exon delta and highest number of ERs with exon delta equal to 0 was chosen as the most accurate transcriptome definition for each tissue. Across all tissues, 50-54% of the ERs tested had an exon delta = 0, suggesting we had defined the majority of ERs accurately. Taking the cerebellum as an example and comparing ER definitions to those which would have been generated applying the default *derfinder* parameters used in the existing literature (MCC: 0.5, MRG: None equivalent to 0), we noted an 96bp refinement in ER size, equating to 67% of median exon size (**Figure 1b & 1c**). In summary, by using known exons to calibrate the detection of transcription, we generated more accurate annotation-agnostic transcriptome definitions for 13 regions of the CNS and a further 28 human tissues.

**Figure 1.**
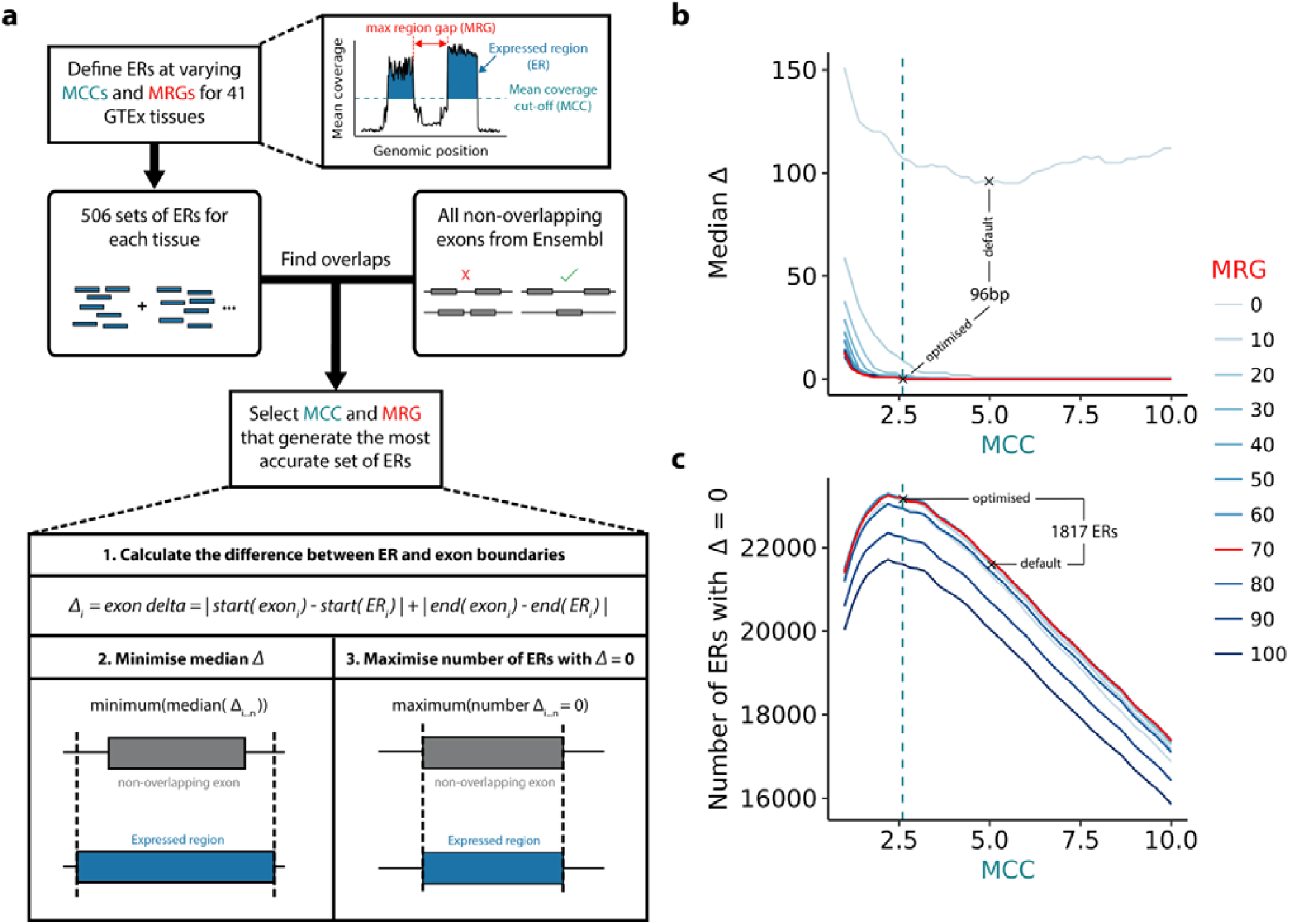
Optimisation of the detection of transcription. **a)** Transcription in the form expressed regions (ERs) was detected in an annotation agnostic manner across 41 human tissues. The mean coverage cut-off (MCC) is the number of reads supporting each base above which that base would be considered transcribed and the max region gap (MRG) is the maximum number of bases between ERs below which adjacent ERs would be merged. MCC and MRG parameters were optimised for each tissue using the non-overlapping exons from Ensembl v92 reference annotation. **b)** Line plot illustrating the selection of the MCC and MRG that minimised the difference between ER and exon definitions (median exon delta). **c)** Line plot illustrating the selection of the MCC and MRG that maximised the number of ERs that precisely matched exon definitions (exon delta = 0). The cerebellum tissue is plotted for (b) and (c), which is representative of the other GTEx tissues. Green and red lines indicate the optimal MCC (2.6) and MRG (70), respectively.

### Novel transcription is most commonly observed in the central nervous system

To assess how much of the detected transcription was novel, we calculated the total size in base pairs of ERs that did not overlap known annotation. ERs were then categorised with respect to the genomic features with which they overlapped as defined by the Ensembl v92 reference annotation (exons, introns, intergenic; **Supplementary Figure 1a**). Those that solely overlapped intronic or intergenic regions were classified as novel. We discovered 8.4 to 22Mb of potentially novel transcription across all tissues, consistent with previous reports that annotation remains incomplete^13,14^. Novel ERs predominantly fell into intragenic regions suggesting that we were preferentially improving the annotation of known genes, rather than identifying new genes (**Figure 2a**). Although novel transcription was found to be ubiquitous across tissues, the abundance varied greatly between tissues (**Figure 2b, 2d, 2e**). To investigate this further, we calculated the coefficient of variation for exonic, intronic and intergenic ERs. We found that the levels of novel transcription varied 3.4-7.7x more between tissues than the expression of exonic ERs (coefficient of variation of exonic ERs: 0.066Mb, intronic ERs: 0.222Mb, intergenic ERs: 0.481Mb). Furthermore, focusing on a subset of novel ERs for which we could infer the precise boundaries of the presumed novel exon (using intersecting split reads), we found that more than half of these ERs were detected in only 1 tissue and that 86.3% were found in less than 5 tissues (**Supplementary Figure 2a**). Even when restricting to ERs derived from only the 13 CNS tissues, 34.3% were specific to 1 CNS region (**Supplementary Figure 2b**). This suggests that novel ERs are largely derived from tissue-specific transcription, potentially explaining why they had not already been discovered.

**Figure 2.**
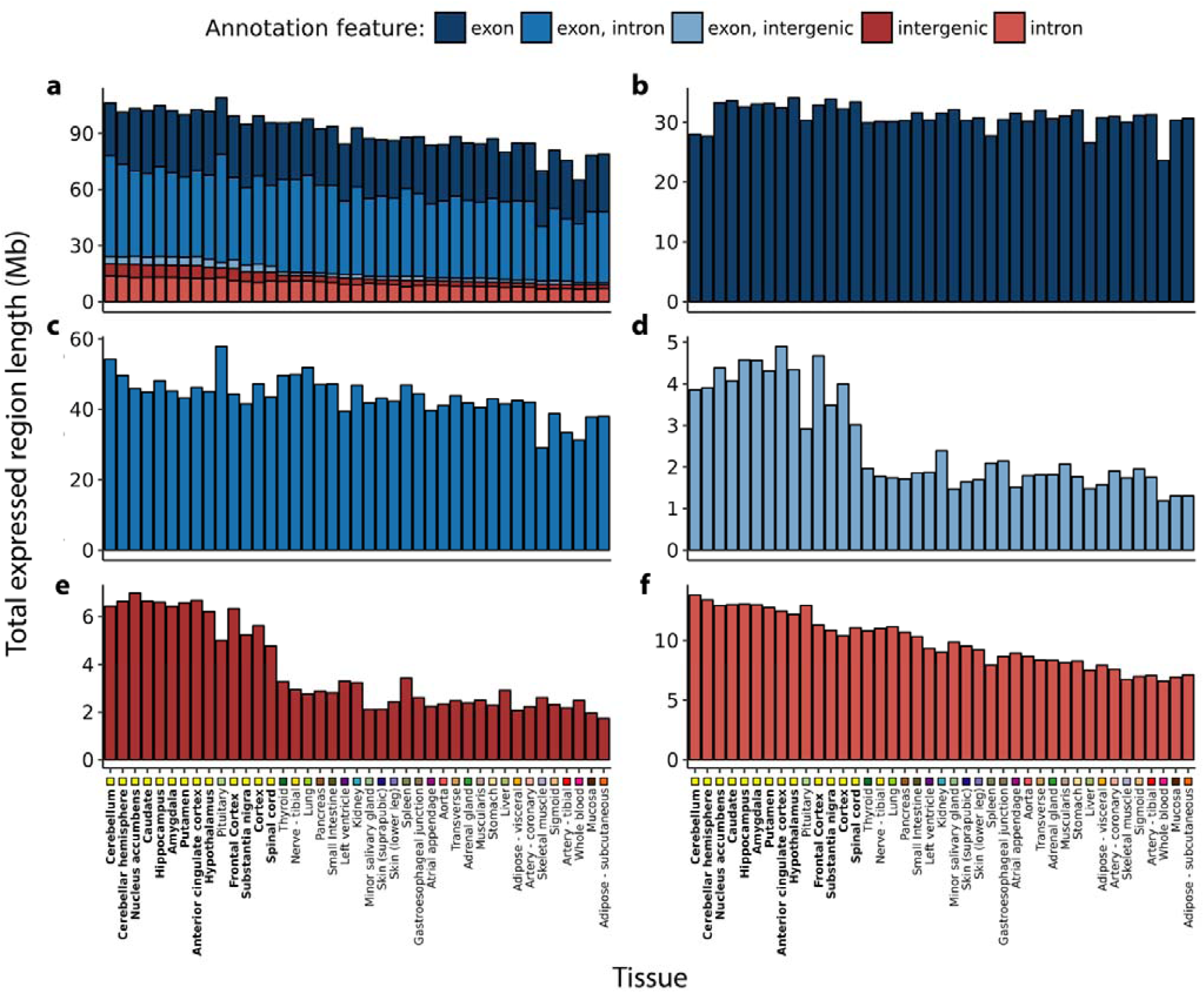
Transcription detected across 41 GTEx tissues categorised by annotation feature. Within each tissue the length of the ERs Mb overlapping **a)** all annotation features **b)** purely exons **c)** exons and introns **d)** exons and intergenic regions **e)** purely intergenic regions **f)** purely introns according to Ensembl v92 was computed. Tissues are plotted in descending order based on the respective total size of intronic and intergenic regions. Tissues are colour-coded as indicated in the x-axis, with GTEx brain regions highlighted with bold font. At least 8.4Mb of novel transcription was discovered in each tissue, with the greatest quantity found within brain tissues (mean across brain tissues: 18.6Mb, non-brain: 11.2Mb, two-sided Wilcoxon rank sum test p-value: 2.35e-10

This finding lead us to hypothesise that genes highly expressed in brain would be amongst the most likely to be re-annotated due to the difficulty of sampling human brain tissue, the cellular heterogeneity of this tissue and the particularly high prevalence of alternative splicing^3^. As we predicted, the quantity of novel transcription found within brain was significantly higher than non-brain tissues (p-value: 2.35e-10) (**Figure 2e & 2f**). In fact, ranking the tissues by descending Mb of novel transcription demonstrated that tissues of the CNS constituted 13 of the top 14 tissues. Interestingly, the importance of improving annotation in the human brain tissue was most apparent when considering purely intergenic ERs and ERs that overlapped exons and extended into intergenic regions (**Figure 2d & 2e**).

This observation raised the question of whether there were specific genic features, which could be used to predict which genes were most likely to be re-annotated (connected to a novel ER). We used logistic regression to determine whether specific properties, including measures of structural gene complexity and specificity of expression to brain increased the likelihood of re-annotation. We also accounted for factors which might be expected to contribute to errors in ER identification, including whether the gene overlapped with another known gene making attribution of reads more complex. We found that the annotation of brain-specific genes and those with higher transcript complexity were more likely to have evidence for incomplete annotation (**Table 1**). Importantly, overlapping genes were not significantly more likely to be re-annotated (taking into account gene length), demonstrating that novel transcription is not merely a product of noise from intersecting genes. Taken together these findings demonstrated that widespread novel transcription is found across all human tissues, the quantity of which varies extensively between tissues. CNS tissues displayed the greatest quantity of novel transcription and accordingly, genes highly expressed in the human brain are most likely to be re-annotated.

**Table 1.** Gene properties influencing re-annotation. Gene characteristics such as brain specificity, transcript count, gene length, mean TPM and whether the gene overlapped with another were used to assess which genes were the most likely to be identified as re-annotated. Brain-specific, longer, protein-coding genes of high transcript complexity were the most likely to be re-annotated. Blue and red highlights positive and negative significant estimates, respectively.

| Gene property | Estimate | P-value |
| --- | --- | --- |
| Brain-specific | 0.093 | *** |
| Transcript count | 0.016 | *** |
| Gene length | 4.18E-07 | *** |
| Gene biotype - protein coding | 0.218 | *** |
| Gene biotype - lincRNA | -0.039 | *** |
| Gene biotype - processed pseudogene | -0.154 | *** |
| Gene biotype - unprocessed pseudogene | -0.093 | *** |
| Gene biotype - other | -0.113 | *** |
| Gene TPM | -2.62E-06 | 0.4 |
| Overlapping gene | 1 | 0.83 |

\*\*\* p <= 2e-16

### Validation of novel transcription across Ensembl versions and within an independent dataset

We recognise that a proportion of novel transcription may originate from technical variability or pre-mRNA contamination. Therefore, we assessed the reliability of novel ERs by classifying ERs using different versions of Ensembl and through an independent dataset. Firstly, we measured how many Kb of the transcription we detected would have been classified as novel with respect to Ensembl v87, but was now annotated in Ensembl v92 and found that across all tissues an average of 68Kb (43-127Kb) had changed status. This value was 5.3x (3.2-10.1x) greater in every tissue compared to the Kb of ERs overlapping exons in Ensembl v87 that had become purely intronic or intergenic in Ensembl v92 (**Figure 3a**). To further assess whether this was greater than what would be expected by chance, we compared the total Kb of novel ERs entering v92 annotation for each tissue to 10,000 sets of random length-matched intronic and intergenic regions. For all tissues, the total Kb of both intronic and intergenic ERs that were now annotated in Ensembl v92 was significantly higher than the total Kb distribution of the randomised negative control regions, implying a high validation rate of novel ERs (**Supplementary figure 3**). Notably, brain regions had significantly higher Kb of ERs entering Ensembl v92 annotation from Ensembl v87 than non-brain tissues, even when subtracting the Kb of ERs leaving Ensembl v87 (p-value: 7.6e-9), suggesting the greater abundance of brain-specific novel transcription was not purely attributed to increased transcriptional noise.

**Figure 3.**
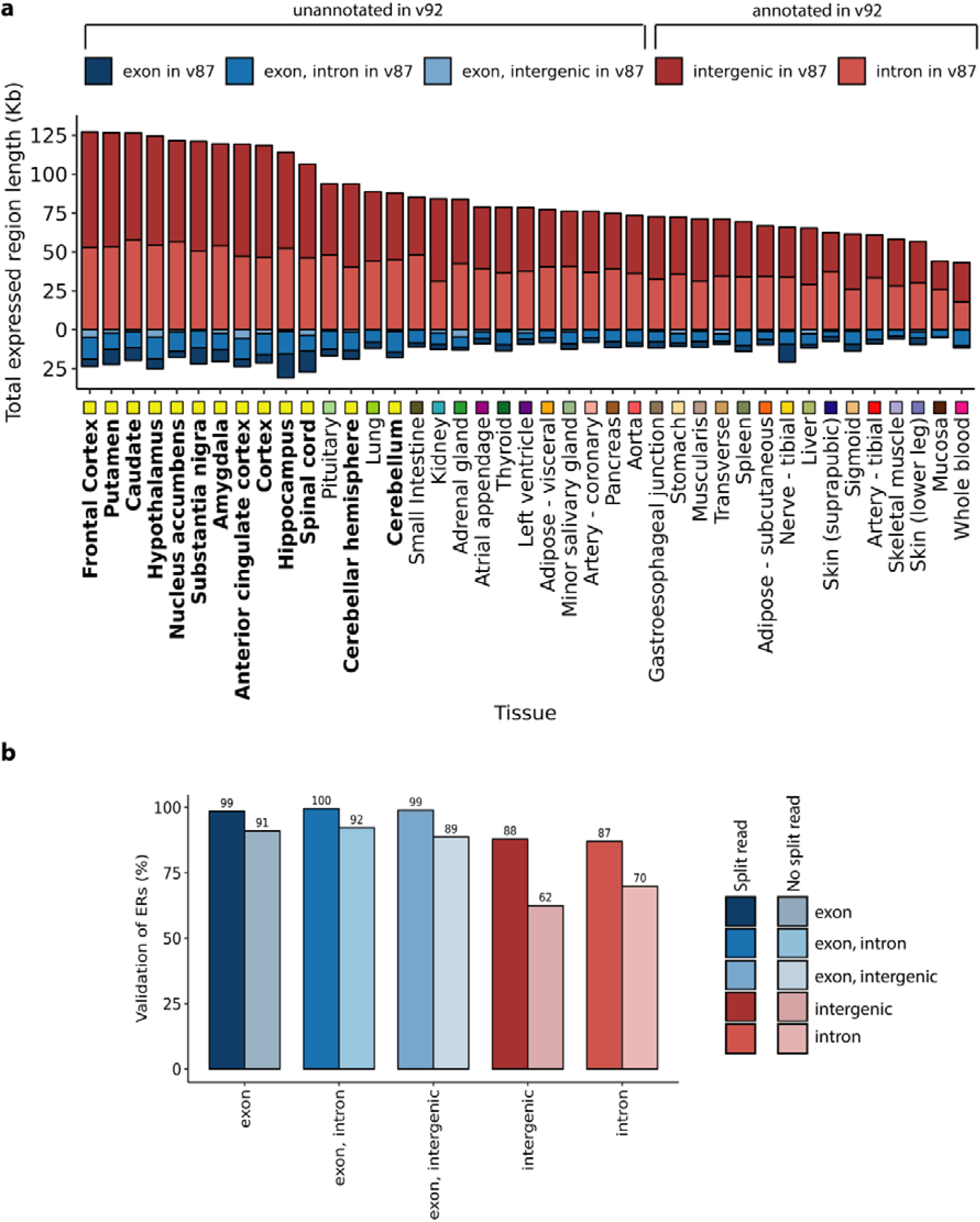
Validation of novel transcription. **a)** The classification of ERs based on v87 and v92 of Ensembl was compared. Across all tissues, the number of intron or intergenic ERs with respect to v87 that were known to be exonic in Ensembl v92 was greater than the number of ERs overlapping exons according to v87 that were now unannotated in v92. Tissues are plotted in descending order based on the total Mb of novel ERs with respect to Ensembl v87 that were validated (classified as exonic in the Ensembl v92). Tissues are colour-coded as indicated in the x-axis, with GTEx brain regions highlighted with bold font. **b)** Barplot represents the percentage of ERs seeding from the GTEx frontal cortex that validated in an independent frontal cortex RNA-seq dataset. ERs defined in the seed tissue were re-quantified using coverage from the validation dataset, after which the optimised mean coverage cut off was applied to determine validated ERs. Colours represent the different annotation features that the ERs overlapped and the shade indicates whether the ER was supported by split read(s).

While our analysis of novel ERs across different Ensembl versions provided a high level of confidence in the quality of ER calling, it was limited to ERs which had already been incorporated into annotation and did not provide an overall indication of the rate of validation across all ERs. Therefore, we investigated whether our GTEx frontal cortex derived ERs could also be discovered in an independent frontal cortex dataset reported by Labadord and colleagues^15^. As expected, ERs which overlapped with annotated exons had near complete validation (>= 89%), but importantly 62% of intergenic and 70% of intronic ERs respectively were also detected in the second independent frontal cortex dataset (**Figure 3b**). While this high validation rate implied the majority of all ERs were reliably detected, we investigated whether a subset of ERs supported with evidence of RNA splicing as well as transcription would have even better rates of validation. Evidence of transcription is provided by the coverage data derived using *derfinder*, whilst split reads, which are reads with a gapped alignment to the genome provide evidence of the splicing out of an intron (**Supplementary figure 1b**). With this in mind, we focused our attention on the putative spliced ERs as indicated by the presence of an overlapping split read. Consistent with expectation, we found that ERs with split read support had higher validation rates than ERs lacking this additional feature. This increase in validation rate for ERs with split read support was greatest for intergenic and intronic ERs with the validation rate rising to 87% for intergenic ERs and 88% for intronic ERs (as compared to 99% for ERs overlapping exons, **Figure 3b**). Even when considering this set of highly validated ERs with split read support, 1.7-3.8Mb of intronic and 0.5-2.2Mb of intergenic transcription was detected across all 41 tissues. Thus, in summary, the majority of novel ERs were reliably detected and validated in an independent dataset.

### Unannotated expressed regions are depleted for genetic variation and some have the potential to be protein coding suggesting they are functionally significant

Given recent reports suggesting widespread transcriptional noise and acknowledging that transcription, even when tissue-specific, does not necessarily translate to function we investigated whether novel ERs were likely to be of functional significance using measures of both conservation and genetic constraint^14,16^. The degree to which a base is evolutionarily conserved across species is strongly dependent on its functional importance and accordingly, conservation scores have been used to aid exon identification^17^. However, this measure is unable to capture genomic regions of human-specific importance. Thus, we investigated novel ERs not only in terms of conservation but also genetic constraint. Constraint scores, measured here as a context-dependent tolerance score (CDTS), represent the likelihood a base is mutated within humans^18^. By comparing our detected novel ERs to 10,000 randomised sets of length-matched intronic and intergenic regions, we found that both intronic and intergenic ERs were significantly less conserved, but more constrained than expected by chance (p-value < 2e-16, **Figure 4a**). This would suggest that they have an important functional role specifically in humans. Furthermore, considering the importance of higher-order cognitive functions in differentiating humans from other species, we measured the constraint of brain-specific novel ERs separately on the basis that these ERs may be the most genetically constrained of all novel ERs identified. Indeed, we found that brain-specific novel ERs were even more constrained than other novel ERs, supporting the view that improvements in gene annotation are likely to have a disproportionate impact on our understanding of human brain diseases.

**Figure 4.**
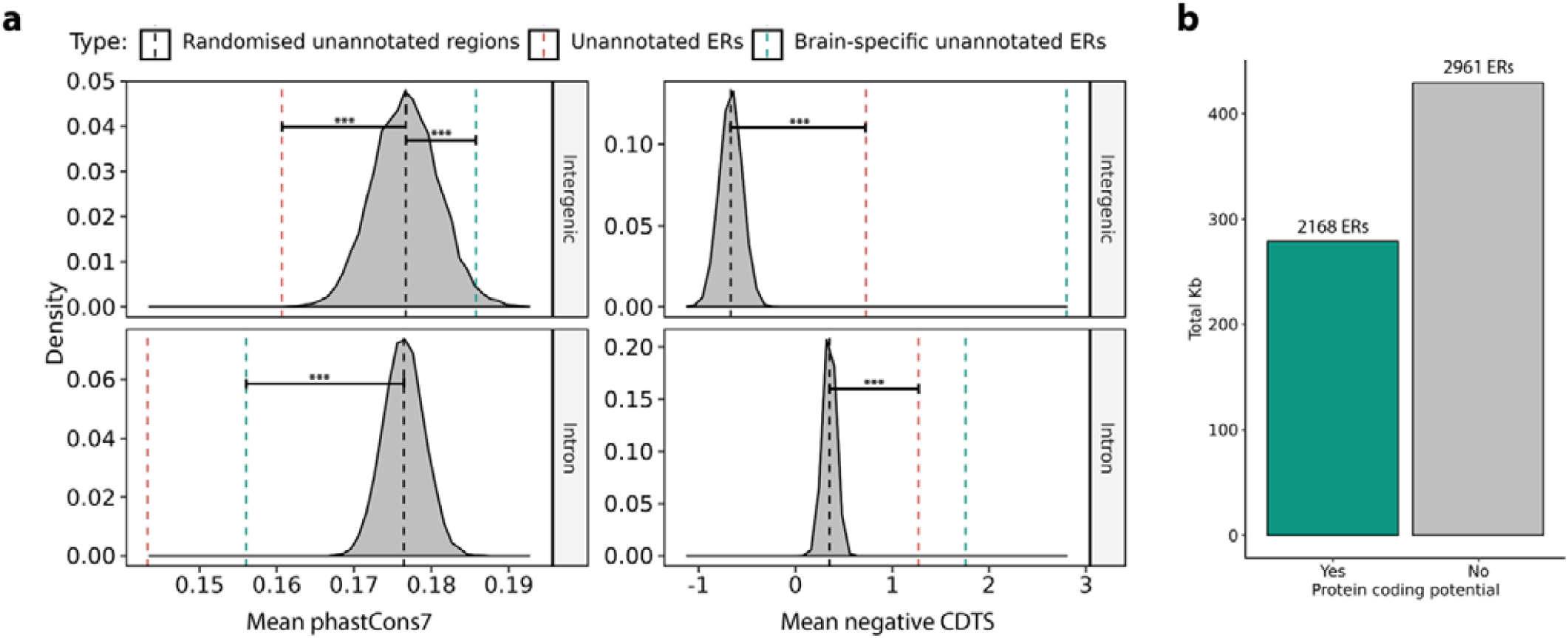
Novel ERs collectively serve an important function for humans and a proportion can form potentially protein coding transcripts. **a)** Comparison of conservation (phastCons7) and constraint (CDTS) of intronic and intergenic ERs to 10,000 sets of random, length-matched intronic and intergenic regions. Novel ERs marked by the red, dashed line are less conserved than expected by chance, but are more constrained. Brain–specific ERs marked by the green, dashed lines are amongst the most constrained. Data for the cerebellum shown and is representative of other GTEx tissues. **b)** The DNA sequence for ERs overlapping 2 split reads was obtained and converted to amino acid sequence for all 3 possible frames. 2,168 ERs (57%) lacked a stop codon in at least 1 frame and were considered potentially protein-coding.

Another metric of functional importance is whether a region of the genome is translated into protein and notably the vast majority of all known Mendelian disease mutations fall within protein-coding regions. For this reason, we investigated whether novel ERs could potentially encode for proteins. Here, we focused on the subset of novel ERs which had evidence of splicing, since the overlapping split reads can be used to assign the precise boundaries of ERs, allowing us to confidently retrieve the DNA sequence and corresponding amino acid sequence for each novel ER. A total of 2,961 ERs covering 274Kb was found to be potentially protein coding, which represented 57% of the ERs analysed (**Figure 4b**). Amongst this set of ERs with protein coding potential, 758 ERs also fell within the top 20% of most constrained regions of the genome. These ERs connect to 694 genes, 30% of which are expressed specifically in the CNS (**Supplementary table 1**). Overall, we discovered that novel ERs broadly are likely to have a human-specific function. We also identified an important subset of novel ERs that have protein coding potential and are highly depleted for genetic variation in humans. Together, this suggested that at least a proportion of novel ERs are functionally significant.

### Incomplete annotation of brain-specific genes may be limiting our understanding of specific CNS-relevant cell types and complex diseases

Given that we discovered the greatest abundance of novel transcription amongst brain tissues, we investigated whether this may be impacting on our understanding of certain cell types within the brain more than others. We tested this by calculating whether our set of 2962 re-annotated brain-specific genes were significantly enriched for cell-type specific genes, when compared to the background list of 2422 brain-specific genes without re-annotations. Of the 13 brain-specific cell types considered, genes specifically expressed by oligodendrocytes had the largest difference in enrichment (p-value of re-annotated: <2e-16; not re-annotated: 0.169), suggesting incomplete annotation was disproportionately limiting our understanding of this cell type (**Figure 5a**). For example, we found that *MBP*, which encodes for myelin basic protein, was amongst those genes re-annotated and with an oligodendrocyte-specific expression profile (**Supplementary figure 4**). In fact, we detected a 48bp ER specific to cortex and striatal tissues (anterior cingulate cortex, cortex, frontal cortex, nucleus accumbens, putamen), which was connected to two flanking protein-coding exons of *MBP*. The ER itself had protein-coding potential and evidence of functional importance specifically in humans, as demonstrated by low mammalian sequence conservation but depletion of genetic variation within humans (phasCons7: 0.03, top 20% CDTS) (Figure 5b). This finding is interesting because *MBP* specifically and oligodendroglial dysfunction more generally, have been implicated in a number of neurodegenerative disorders, including multiple system atrophy, which is characterised by myelin loss and degeneration of striatum and cortical regions^19^, as well as schizophrenia and Parkinson’s disease^19–21^.

**Figure 5.**
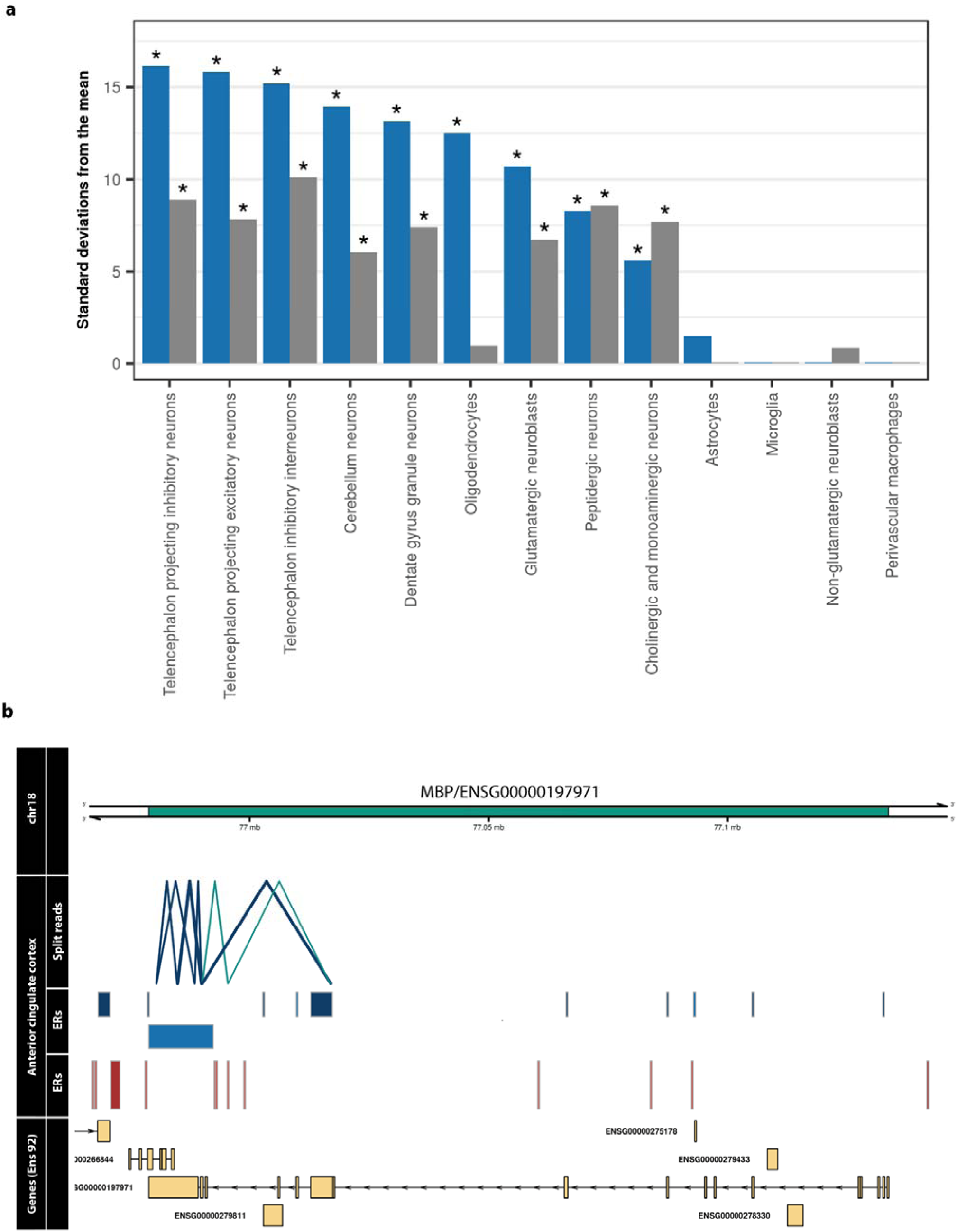
Incomplete annotation of genes disproportionately affects oligodendrocytes. a) Bar plot displaying the enrichment of re-annotated and not re-annotated genes within brain cell-type specific gene sets. Blue bars represent the re-annotated genes and grey are those without re-annotations. Of all analysed cell-types, the greatest difference between enrichment of re-annotated and not re-annotated was observed in oligodendrocytes. b) Novel potentially protein coding ER discovered in MBP, with an oligodendrocyte specific expression

These observations led us to postulate whether incomplete annotation could also be hindering our understanding of complex neurodegenerative and neuropsychiatric disorders. Therefore, we assessed whether our list of re-annotated genes was enriched for genes associated with complex forms of neurodegenerative, neuropsychiatric or other neurological conditions. This analysis was performed by using the Systematic Target OPportunity assessment by Genetic Association Predictions (STOPGAP) database, which provides an extensive catalogue of human genetic associations mapped to effector gene candidates (see detailed methods)^22^. Interestingly, we found that genes associated with neurodegenerative disorders were significantly over-represented within our re-annotated set (p-value: 0.004, **Supplementary Table 2**). In particular, important neurodegenerative disease genes such as *SNCA, APOE* and *CLU* were amongst those re-annotated, suggesting that despite being extensively studied the annotation of these genes remains incomplete (complete list found in **Supplementary Table 3**). Thus, we demonstrate that incomplete annotation of brain-specific genes may be hindering our understanding of specific cell types and complex neurodegenerative disorders.

### Incomplete annotation of OMIM genes may limit genetic diagnosis, particularly for neurogenetic disorders

Since re-annotation of genes already known to cause Mendelian disease would have a direct impact on clinical diagnostic pipelines, we specifically assessed this gene set. Novel ERs were first connected to known genes using split reads (**Supplementary figure 1b**). Next, we filtered for OMIM-morbid genes and found that 63% of this set of OMIM-morbid genes were re-annotated and 14% were connected to a potentially protein-coding ER, suggesting that despite many of these genes having been extensively studied, the annotation of many OMIM-morbid genes remains incomplete (**Figure 6a**). Given that OMIM-morbid genes often produce abnormalities specific to a given set of organs or systems, we investigated the relevance of novel transcription to disease by matching the human phenotype ontology (HPO) terms obtained from the disease corresponding to the OMIM-morbid gene, to the GTEx tissue from which ERs connected to that gene were derived. We discovered that 72% of re-annotated OMIM-morbid genes had an associated novel ER originating from a phenotypically relevant tissue (**Figure 6b**). This phenomenon was exemplified by the OMIM-morbid gene *ERLIN1*, which when disrupted is known to cause spastic paraplegia 62 (SPG62), an autosomal recessive form of spastic paraplegia, which has been reported in some families to cause not only lower limb spasticity, but also cerebellar abnormalities^23^. We detected a cerebellar-specific novel ER that was intronic with respect to ERLIN1. The novel ER had the potential to code for a non-truncated protein and connected through intersecting split reads to two flanking, protein-coding exons of *ERLIN1*, supporting the possibility of this ER being a novel protein-coding exon. Furthermore, the putative novel exon was highly conserved (phastcons7 score: 1) and was amongst the top 30% most constrained regions in the genome, suggesting it is functionally important both across mammals and within humans (**Figure 6c**).

**Figure 6.**
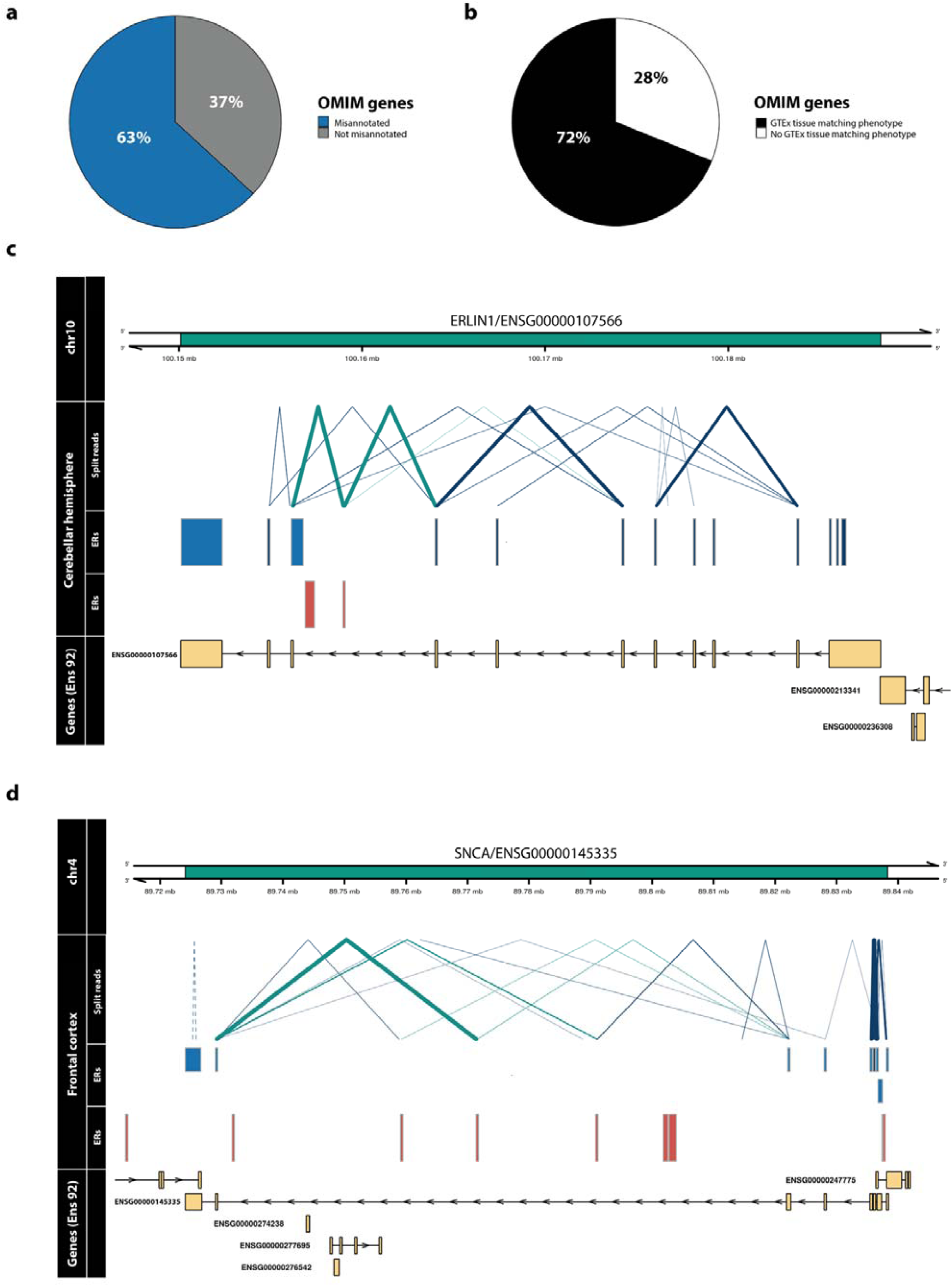
Re-annotation of OMIM genes. **a)** A novel ER connected through a split read was discovered for 63% of OMIM-morbid genes. **b)** Comparison of the phenotype (HPO terms) associated with each re-annotated OMIM-morbid gene and the GTEx tissue from which novel ERs were derived. Through manual inspection, HPO terms were matched to disease-relevant GTEx tissues and for 72% of re-annotated OMIM the split reads and ERs overlapping the genomic region derived from the labelled tissue. Blue ERs overlap known exonic regions and red ERs fall within intronic or intergenic regions. Blue split reads overlap blue ERs, while green split reads overlap both red and blue ERs, connecting novel ERs to OMIM-morbid genes. Thickness of split reads represents the proportion of samples of that tissue in which the split read was detected. Only partially annotated split reads (solid lines) and unannotated split reads (dashed lines) are plotted. The last track displays the genes within the region according to Ensembl v92, with all known exons of the gene collapsed into one “meta” transcript.

Similarly, we detected a brain-specific novel ER in the long intron of the gene *SNCA*, which encodes alpha-synuclein protein implicated in the pathogenesis of Mendelian and complex Parkinson’s disease. This ER connected to two flanking protein-coding exons through split reads (**Figure 6d**) and appeared to also have coding potential. Interestingly, while the ER sequence is not conserved within mammals (phastcons7 score: 0.09) or primates (phastcons20 score: 0.21), it is in the top 19% of most constrained regions in the genome suggesting it is of functional importance specifically in humans. We validated the existence of this ER both in silico and experimentally. The expression of this ER was confirmed in silico using an independent frontal cortex dataset reported by Labadord and colleagues^15^. Using Sanger sequencing, we validated the junctions intersecting the ER and the flanking exons in RNA samples originating from pooled human frontal cortex samples (**Supplementary Figure 5**). In order to gain more information about the transcript structure in which the novel ER was contained, we also performed Sanger sequencing from the first (ENSE00000970013) and last coding exons (ENSE00000970014) of *SNCA* to the novel ER. This implied a full transcript structure containing a minimum of 609bp with the novel ER predicted to add an additional 63 amino acids (45% of existing transcript size). This example highlights the potential of incomplete annotation to both hinder genetic diagnosis and limit our understanding of a common complex neurological disease. Variants located in the novel ER linked to *SNCA* would not be captured using using whole exome sequencing (WES) and if identified in WGS or through GWAS would be misassigned as non-coding variants.

## Discussion

In this study, we demonstrate that novel transcription though commonly detected across all tissues, disproportionately affects genes highly expressed in the brain and some brain-specific cell types, namely oligodendrocytes. We provide evidence to suggest that novel ERs are functionally important, since they are more depleted for genetic variation within humans than would be expected by chance and some have the potential to code for protein. Furthermore, we find that genes known to cause Mendelian and complex neurodegenerative disorders are enriched amongst the set of genes we reannotate. In order to illustrate the potential impact of incomplete annotation on such disorders, we highlight the specific example of *SNCA*, a gene implicated in Mendelian and complex Parkinson’s disease. We experimentally validate the existence of a novel transcript of *SNCA* containing a potentially protein coding novel ER. Together, this suggests that incomplete annotation may be limiting our understanding of both Mendelian and common complex diseases.

We find that the majority of probable novel exons we detect have a restricted expression pattern across tissues. The practical difficulty of accessing the brain reduces the number of available brain-specific datasets and its regional and cellular heterogeneity is one of the factors driving the high number of brain-specific transcripts. Furthermore, since our approach does not depend on conservation across species to annotate novel exons, we are able to identify ERs which are likely to be of human-specific importance^18^. In fact, we find that brain-specific ERs have the highest constraint scores, emphasising their specific importance in humans. Together these factors suggest that the resource we have generated will have the greatest impact on neurogenetic disorders.

Finally, we release our results through a dedicated web resource, vizER (http://rytenlab.com/browser/app/vizER), which enables individual genes to be queried for incomplete annotation as well as the download of all novel ER definitions. We believe that vizER will be an important resource for clinical scientists in the diagnosis of Mendelian disorders, neuroscientists studying individual gene structures and functions, and with the emergence of larger long read sequencing data sets will accelerate novel transcript discovery particularly in human brain.

## Online Methods

### OMIM data

Phenotype relationships and clinical synopses of all Online Mendelian Inheritance in Man (OMIM) genes were downloaded using http://api.omim.org on the 29^th^ of May 2018^24^. OMIM genes were filtered to exclude provisional, non-disease and susceptibility phenotypes retaining 2,898 unique genes that were confidently associated to 4,034 Mendelian diseases. Phenotypic abnormality groups were linked to corresponding affected Genotype-Tissue Expression (GTEx) tissues through manual inspection of the HPO terms within each group by a medical specialist^11^.

### GTEx data

RNA-seq data in base-level coverage format for 7,595 samples originating from 41 different GTEx tissues was downloaded using the R package recount version 1.4.6^4^. Cell lines, sex-specific tissues and tissues with 10 samples or below were removed. Samples with large chromosomal deletions and duplications or large CNVs previously associated with disease were filtered out (smafrze = “USE ME”). Coverage for all remaining samples was normalised to a target library size of 40 million 100bp reads using the area under coverage value provided by recount2. For each tissue, base-level coverage was averaged across all samples to calculate the mean base-level coverage. GTEx split read data, defined as reads with a non-contiguous gapped alignment to the genome, was downloaded using the recount2 resource and filtered to include only split reads detected in at least 5% of samples for a given tissue and those that had available donor and acceptor splice sequences.

### Optimising the detection of transcription

Transcription was detected across 41 GTEx tissues using the package derfinder version 1.14.0^12^. The mean coverage cut-off (MCC), defined as the number of reads supporting each base above which bases were considered to be transcribed, and max region gap (MRG), defined as the maximum number of bases between expressed regions (ERs) below which adjacent ERs will be merged, were optimised. Optimisation was performed using 156,674 non-overlapping exons (defined by Ensembl v92) as the gold standard^10^. Exon biotypes of all Ensembl v92 exons were compared to this set of non-overlapping exons to ensure we were not preferentially optimising for one particular biotype (**Supplementary figure 6**). Non-overlapping exons were selected as these definitions would be least likely to be influenced by ambiguous reads. For each tissue, we generated ERs using mean coverage cut-offs increasing from 1 to 10 in steps of 0.2 (46 cut-offs) and max gaps increasing from 0 to 100 in steps of 10 (11 max region gaps) to produce a total of 506 unique transcriptomes. For each set of ERs, we found all ERs that intersected with non-overlapping exons, then calculated the exon delta by summing the absolute difference between the start/stop positions of each ER and the overlapping exon (Figure 1a). Situations in which a single ER overlapped with multiple exons were removed to avoid assigning the ER to an incorrect exon when calculating downstream optimisation metrics. For each tissue, we selected the mean coverage cut-off and max region gap, which minimised the difference between ER and “gold standard” exon definitions (median exon delta) and maximised the number of ERs that precisely matched the boundaries of exons (number of ERs with an exon delta equal to 0). All ERs that were <3bp in width were removed as these were below the minimum size of a microexon^25^.

### Calculating the transcriptome size per annotation feature

ERs were classified with respect to the annotation feature (exon, intron, intergenic) with which they overlapped. A minimum of 1bp overlap was required for an ER to be categorised as belonging to a given annotation feature. ERs overlapping multiple annotation features were labelled with a combination of each. This generated 6 distinct categories – “exon”, “exon, intron”, “exon, intergenic”, “exon, intergenic, intron”, “intergenic” and “intron” (**Supplementary figure 1a**). ERs classified as “exon, intergenic, intron” were removed from all downstream analysis as these formed only 0.54% of all ERs and were presumed to be technical artefacts generated from regions of dense, overlapping gene expression. For each tissue, the length of all ERs within each annotation feature was summed generating the total Mb of ERs per annotation feature. Normalised variance of exonic, intronic and intergenic ERs was calculated by dividing the standard deviation of the total Mb of ERs across tissues by the mean total Mb of ERs for each annotation feature. To compare between brain and non-brain tissues, the total Mb of intronic and intergenic ERs were first summed together to generate an overall measure of novel transcription abundance across brain and non-brain tissues, then a two-sided Wilcoxon rank sum test was applied.

### Annotating ERs with split read data

Intronic and intergenic ERs were connected to known genes using reads, which we term split reads, with a gapped alignment to the genome, presumed to be reads spanning exon-exon junctions (Supplementary figure 2b). Such exon-exon junctions are defined as non-contiguous reads which fall on the boundary between two exons of the same mRNA molecule, therefore when aligned to the genome these reads have a break in the middle indicating the splicing out of an intron. Split read data was categorised into three groups: annotated split reads, with both ends falling within known exons; partially annotated split reads, with only one end falling within a known exon; and unannotated split reads, with both ends within intron or intergenic regions. In this way, intron and intergenic ERs that overlapped with partially annotated split reads were connected to known genes.

### Validation of detected transcription

Transcription was validated across different versions of Ensembl and within an independent dataset. ERs that overlapped purely intronic or intergenic regions according to Ensembl v87, but fell within exons according to v92, were counted as novel transcription that was validated in later versions of Ensembl. Furthermore, ERs overlapping exonic regions in Ensembl v87 now classified as intronic or intergenic in v92 were measured to control for expected corrections in gene definitions. To assess whether the total Kb of validated novel ERs entering v92 annotation was greater than what would be expected by chance, we generated 10,000 random sets of length-matched regions for each tissue that were intronic or intergenic with respect to Ensembl. Using a one sample Wilcoxon test, we compared the total Kb of intronic and intergenic ERs entering annotation to the total Kb distribution of the randomised intronic and intergenic regions, respectively.

Validation within an independent dataset was performed using RNA-seq coverage data from 49 control frontal cortex (BA9) samples originally reported by Labadorf and colleagues (2015) and available via the recount R package version 1.4.6^4,15^. ERs derived from the GTEx frontal cortex (BA9) data were re-quantified using this independent frontal cortex dataset and those that had a mean coverage of at least 1.4 (the optimised MCC for the GTEx frontal cortex data), were counted as novel transcription that was validated.

### Analysing the conservation and constraint of novel ERs

Conservation scores in the form of phastCons7 (derived from genome-wide alignments of 7 mammalian species) were downloaded from UCSC^26,27^. Constraint scores generated from the genome-wide alignment of 7,794 unrelated human genomes were downloaded as context dependent tolerance scores (CDTS)^18^. The raw phastCons7 and CDTS were in bins of 1bp and 10bp, respectively, therefore when annotating the corresponding positions of ERs, we aggregated each score as a mean across the entire genomic region of interest. To account for missing CDTS values, we calculated the coverage of each ER by dividing the number of bases annotated by the CDTS by the total length of the ER. For all downstream analysis, we filtered out ERs for which CDTS coverage was less than 80%.

To assess whether our novel ERs were more constrained or conserved than by expected by chance, we compared the phastCons7 and CDTS of novel ERs to 10,000 randomised length-matched sets of intronic and intergenic ERs for each tissue. For each of the 10,000 iterations, we first selected a random intronic or intergenic region that was larger than the respective ER, then selected a random segment along the randomised region which matched the length of the corresponding ER. The randomised regions were annotated with constraint scores and CDTS using the aforementioned method. The mean CDTS and phastCons7 of the novel ERs (split by annotation feature) were compared to the corresponding distribution of CDTS and phastCons7 of the randomised regions using a one sample, two-tailed t-test. For easier interpretation when plotting, CDTS scores have been converted to their opposite sign, therefore for both phastCons and CDTS, the higher the value the greater the magnitude of conservation or constraint as shown in Figure 4a.

### Checking ER protein coding potential

Intronic and intergenic ERs that were intersected by 2 split reads were extracted. The split reads were used to determine the precise boundaries of the ER. The R package Biostrings version 2.46.0 was used to extract the DNA sequence corresponding to the ER genetic co-ordinates from the genome build hg38^28^. Since the translation frame was ambiguous without knowledge of the other exons that are part of the transcript that included the novel ER, we converted the DNA sequence to amino acid sequence for all three possible frames starting from the first, second or third base. Any ER that had at least 1 frame that did not include a stop codon was considered to be potentially protein coding.

### Gene properties influencing re-annotation

All Ensembl v92 genes were marked with a 1 or a 0 depending on whether we detected a re-annotation for that gene in the form of an ER connected to the gene using a split read, with 1 representing a detected re-annotation event. Details of gene length, biotype, transcript count and whether the gene overlapped another gene were retrieved from the Ensembl v92 database. Brain-specificity was assigned using the Finucane dataset and selecting the top 10% of brain-specific genes when compared to non-brain tissues^29^. Mean gene TPM was calculated by downloading tissue-specific TPM values from the GTEx portal and summarised by calculating the mean across all tissues. The list of OMIM genes (May 2018) was used to assign whether a gene was known to cause disease or not. We used a logistic regression to test whether different gene properties significantly influenced the variability of re-annotation (formula = re-annotated ∼ brain specific + mean TPM + overlapping gene + transcript count + gene biotype + gene length).

### Sanger sequencing of novel junctions

Commercially purchased (Takara) frontal cortex and cerebellum RNA samples, isolated from individuals of European descent, were used for validation of novel junctions detected in *SNCA* and *ERLIN1* respectively. Tissues were chosen to match the tissue in which the re-annotation for each gene was detected. Reverse transcription was performed using 1ug of RNA from each tissue, then converted to cDNA using the High-Capacity cDNA Reverse Transcription Kit with RNase Inhibitor (Applied Biosystems) and random primers as per manufacturer’s instructions. Primers were designed to span predicted exon-exon junctions using Primer-BLAST (NCBI) and ordered from Sigma (**Supplementary Table 4**). PCR was performed using FastStart PCR Master (Roche) and enzymatic clean-up of PCR products was performed using Exonuclease I (Thermo Scientific) and FastAP Thermosensitive Alkaline Phosphatase (Thermo Scientific). Sanger sequencing was performed using the BigDye terminator kit (Applied Biosystems) and sequences were viewed and exported using CodonCode Aligner (V. 8.0.2). Sequences were blatted against the human genome (hg38) and alignment visually inspected for confirmation of validation.

### Expression-weighted cell-type enrichment (EWCE): evaluating enrichment of theta-correlated genes

EWCE was used to determine whether brain-specific genes (both re-annotated and not re-annotated) have higher expression within particular cell types than expected by chance^30^. As our input, we used 1) neuronal and glial clusters of the central nervous system (CNS) identified in the Linnarsson single-cell RNA sequencing dataset (amounting to a subset of 114 of the original 265 clusters identified) and 2) lists of genes split by whether or not they were re-annotated, and if re-annotated, by their overlap with Ensembl v92 annotation features (see **Supplementary Table 5** for full list of CNS neuronal clusters and genes used)^31^. For each gene in the Linnarsson dataset, we estimated its cell-type specificity (the proportion of a gene’s total expression in one cell type compared to all cell types) using the ‘generate.celltype.data’ function of the EWCE package. EWCE with the target list was run with 100,000 bootstrap replicates, which were sampled from a background list of genes that excluded all genes without a 1:1 mouse:human ortholog. We additionally controlled for transcript length and GC-content biases by selecting bootstrap lists with comparable properties to the target list. We performed the analysis with major cell-type classes classes (e.g. “astrocyte”, “microglia”, etc.). Data are displayed as standard deviations from the mean, and any values < 0, which reflect a depletion of expression, are displayed as 0. P-values were corrected for multiple testing using the Benjamini-Hochberg (FDR) method over all cell types and gene lists displayed.

### Enrichment of re-annotated genes for neurological disorder associated genes

The STOPGAP database detailing all genes associated with 4684 GWASs was downloaded from https://github.com/StatGenPRD/STOPGAP/blob/master/STOPGAP_data/stopgap.bestld.RData. To select which genes were associated to a GWAS, the “best gene” as determined by STOPGAP using functional evidence was used^22^. The medical subject heading for each disease was used to further subgroup GWASs into 4 categories; neurodegenerative, neuropsychiatric, other neurological conditions and the remaining as other (**Supplementary table 6**). For each of the subgroups, we generated a contingency table, counting the number of genes that were re-annotated or not in relation to whether they fell into that particular subgroup. For genes that were overlapping between GWASs, we classified a gene to be part of a subgroup if it was associated with at least 1 GWAS contained in that subgroup. A Fisher’s Exact test was used to examine whether our re-annotated gene list was significantly enriched for genes from any of the subgroups. Benjamini-Hochberg (FDR) method was used for to correct for multiple testing.

## Acknowledgements

S.G. was supported through the award of an Alzheimer’s Research UK PhD fellowship. R.H.R. was supported through the award of a Leonard Wolfson Doctoral Training Fellowship in Neurodegeneration. J.H. and M.R. were supported by the UK Medical Research Council (MRC), with J.H. supported by a grant (MR/N026004/) and M.R. through the award of a Tenure-track Clinician Scientist Fellowship (MR/N008324/1). J.H. was also supported by the UK Dementia Research Institute.

## Author contributions

D.Z., S.G. and M.R. conceived and designed the study. D.Z. analysed the data, generated figures and together with M.R. wrote the first draft of the manuscript. R.H.R. performed analysis and generated figures for the cell-type specific section. S.G.R. and J.B. developed and deployed the vizER online platform. Sanger sequence validation was performed by B.C. and W.L. T.C. helped manually associate OMIM phenotypes to GTEx tissues. S.G, L.C.T, J.B., K.D., A.P. and M.R. helped guide and troubleshoot analyses. L.C.T and A.E.F helped with the use of the recount2 data. D.Z., S.G., R.H.R., J.H., L.C.T. and M.R. contributed to the critical analysis of the manuscript.

